# OrthologAL: A Shiny application for quality-aware humanization of non-human pre-clinical high-dimensional gene expression data

**DOI:** 10.1101/2024.11.25.625000

**Authors:** Rishika Chowdary, Robert K. Suter, Matthew D’Antuono, Cynthia Gomes, Joshua Stein, Ki-Bum Lee, Jae K Lee, Nagi G. Ayad

## Abstract

Single-cell and spatial transcriptomics provide unprecedented insight into the inner workings of disease. Pharmacotranscriptomic approaches are powerful tools that leverage gene expression data for drug repurposing and treatment discovery in many diseases. Multiple databases attempt to connect human cellular transcriptional responses to small molecules for use in transcriptome-based drug discovery efforts. However, pre-clinical research often requires *in vivo* experiments in non-human species, which makes capitalizing on such valuable resources difficult. To facilitate the application of pharmacotranscriptomic databases to pre-clinical research models and to facilitate human orthologous conversion of non-human transcriptomes, we introduce OrthologAL. OrthologAL leverages the BioMart database to access different gene sets from Ensembl, facilitating the interaction between these servers without needing user-generated code. Researchers can input their single-cell or other high-dimensional gene expression data from any species, and OrthologAL will output a human ortholog-converted dataset for download and use. To demonstrate the utility of this application, we characterized orthologous conversion in single-cell, single-nuclei, and spatial transcriptomic data derived from common pre-clinical models, including patient-derived orthotopic xenografts of medulloblastoma, and mouse and rat models of spinal cord injury. We show that OrthologAL can convert these data types efficiently to that of corresponding orthologs while preserving the dimensional architecture of the original non-human expression data. OrthologAL will be broadly useful for applying pre-clinical, high-dimensional transcriptomics data in functional small molecule predictions using existing human-annotated databases.

## 1 INTRODUCTION

Orthologous genes are paired genes in different species that originate from a common ancestor and typically have overlapping or similar functions^1^. Computational identification of orthologous genes in expression data is commonly facilitated by The Ensembl BioMart server ^2,3^, which is a free and scalable database supporting large data resources, including Ensembl and UniProt^4,5^. Seurat is a widely used R package designed for the analysis of single-cell RNA-seq (scRNA-seq) data^6–10^. Seurat Objects in R serve as containers for single-cell and spatial transcriptomics data and facilitate accessible single-cell data manipulation and analysis using the Seurat library ^8,11,12^. While Ensembl’s BioMart data-mining tool allows for the extraction of Ensembl data without coding knowledge and provides access to the corresponding R package, BioMart, the process of orthologous conversion of expression matrices or Seurat objects is not trivial for non-coders. In addition, existing approaches do not provide Seurat integration, nor do they quantify cell and pixel-level quality control (QC) metrics from high-dimensional single-cell data in the context of orthologous conversion.

To address these limitations, we created OrthologAL as a user-friendly method of performing human orthologous conversion on Seurat objects while maintaining awareness of quality control metrics before and after conversion. OrthologAL leverages the BioMart server to access gene sets from multiple species and extract them through common identifiers such as Ensembl ID, species gene symbol, and attribute. These common identifiers help us to find genes from various species in their database, and subsequently map them to their corresponding orthologous human genes. OrthologAL’s Shiny^13,14^ web application interface allows seamless integration of the BioMaRt package with Seurat, and provides an interface to explore quality control metrics (**Figure 1**). Uniquely, OrthologAL also facilitates the orthologous conversion and harmonization of input from models containing expression data from multiple species, such as that from xenograft models.

**Figure 1.**
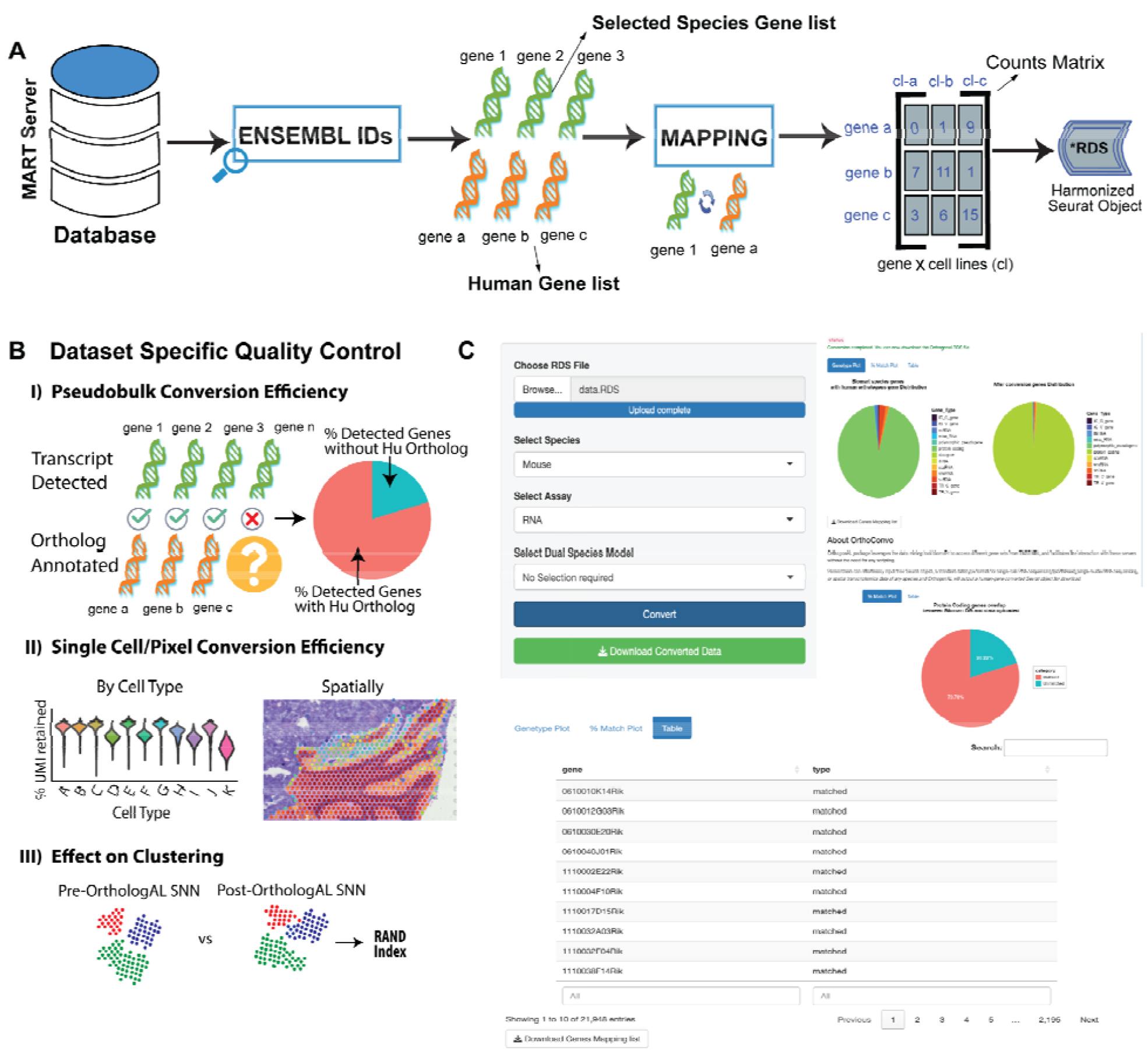
OrthologAL facilitates the conversion of non-human, single-cell, single-nuclei or spatial transcriptomics data to that of orthologous human genes. **A)** Schematic of OrthologAL conversion of non-human gene expression to human ortholog expression utilizing BioMart and Seurat object formats. **B)** Dataset-specific quality controls for OrthologAL-based humanization are assessed using multiple approaches, including pseudobulk conversion efficiency, single-cell and pixel-level conversion efficiency, and the effects on SNN clustering using the RAND index. **C)** Screenshot of the OrthologAL graphical user interface depicting user controls and quality control output.

Single-cell RNA sequencing applications have provided significant insights into diseases and biological processes. Through the humanization of pre-clinical scRNA-seq and spatial transcriptomics (spatialRNA-seq) data, information can be leveraged using human-derived external databases such as the LINCS L1000 dataset, as we and others have previously done with bulk gene expression data^15–19^. We, therefore, sought to explore the utility of OrthologAL in two distinct medical conditions that share benefits from high-dimensional transcriptomic profiling and routinely use pre-clinical animal models in research: central nervous system (CNS) trauma and cancer.

CNS trauma can be clinically devastating and exceptionally difficult to treat, as observed in the case of spinal cord injury (SCI). SCI is typically the result of trauma to the vertebrae or surrounding structures causing contusion or laceration of the spinal cord. The response is orchestrated by a myriad of discrete cell types ^20,21^. Medulloblastoma (MB) is a pediatric embryonal tumor that arises in the cerebellum and can be classified into 4 main molecular subgroups: Sonic Hedgehog (SHH), Wingless (WNT), Group 3 (G3) and Group 4 (G4)^22,23^. MB resolved at the single-cell level reveals transcriptional states spanning neurodevelopmentally-rooted trajectories.^24^ Both MB and SCI require careful pre-clinical study and the use of animal models to develop novel therapies, rehabilitation, and treatment strategies. The analysis of single-cell and spatial transcriptomics have significantly advanced our understanding of both conditions, yet there are currently no simple, “one-click” methods for orthologous conversion of such datasets.

To demonstrate the utility of OrthologAL, we applied it to scRNA-seq and spatialRNA-seq datasets from genetically engineered mouse models (GEMMs) and patient-derived orthotopic xenograft (PDOX) models of MB. Additionally, we converted single-cell and single-nuclei RNA sequencing (snRNA-seq) data from both mouse and rat models of SCI to human orthologs using OrthologAL. With these use cases, we show that OrthologAL-mediated orthologous conversion preserves the structure and content of high-dimensional transcriptomic data for each species and data type presented. We anticipate that OrthologAL will have wide-ranging applications in various pre-clinical studies across multiple disease areas, beyond MB and SCI, with specific utility in pharmacotranscriptomics and drug discovery.

## 2 METHODS

### 2.1 OrthologAL user interface

The OrthologAL application was developed using Shiny^13^ and enables access to orthologous human gene sets from different species via BioMart without coding (**Figure 1**). OrthologAL takes a Seurat v5 object R data file (.RDS)^11,12^ as input and is backward-compatible with previous versions of Seurat. A drop-down menu allows the user to easily select the desired Seurat object ‘.RDS’ file for local upload to the application. Additional drop-downs allow the user to select the species profiled by their uploaded data, and choose the appropriate assay (i.e. ‘RNA’, ‘SCT’, etc.) of the Seurat object where the desired expression data to be converted are stored. By default, OrthologAL supports the orthologous conversion of mouse and rat expression data. However, additional species can be processed and converted using a custom species input field. The user interface is shown in **Figure 1C**.

### 2.2 OrthologAL pipeline, BioMart integration, and humanized assay storage

Once data is uploaded to OrthologAL and configuration parameters are set, the orthologous conversion of the Seurat object can be initiated with a single click. A status bar will appear to update the user on the progress of the conversion. During conversion, Ensembl IDs are queried against the input species code or official gene symbol for that species if Ensembl IDs are not the primary transcript identifier within the uploaded Seurat object (**Figure 1A**). This process is streamlined as data frames containing matched ortholog data for both rat and mouse genomes are stored within the OrthologAL R package to allow for rapid conversion. OrthologAL-mediated conversion will create a new, converted, assay in the Seurat object with the suffix ‘_ortho’ appended to the original assay name (i.e. ‘RNA_ortho’, ‘Spatial_ortho’, etc.). The row names of this assay are then set as the corresponding orthologous human gene symbols (HGNC (HUGO Gene Nomenclature Committee) symbols), and the associated counts are retained. The converted Seurat .RDS file will then be available to download by clicking the “Download Converted Data” button.

### 2.3 OrthologAL pipeline for humanization and harmonization of dual-species input

In the event a user has a Seurat object containing dual-species expression data, such as that from a patient-derived orthotopic xenograft (PDOX)^25^ in a mouse, they can select that option from the “Select Dual Species Model” drop-down menu. While running, OrthologAL will then recognize the standard format of count matrices output from a CellRanger count alignment run with a dual-species mouse and human transcriptome. These count matrices contain Ensembl IDs from both human and mouse transcriptomes, and OrthologAL will selectively convert the expression data of mouse transcripts to their corresponding human orthologs. OrthologAL will then merge the humanized expression matrix with the original human counts present within the dataset and will store this harmonized data within a new assay slot, as explained above (**Figure 1A**). The converted Seurat .RDS file can then be downloaded as well.

### 2.4 OrthologAL data efficiency quantification

Upon successful conversion of non-human expression to that of human orthologs, OrthologAL will generate QC graphics depicting conversion metrics. These include figures showing the number of genes from the uploaded object that were successfully matched to human orthologs within the BioMart database (DB). Additional graphics depict other QC metrics, including the proportion of the biotypes (transcript classifications, e.g., protein-coding of overall orthologous genes for the input species, the proportion of biotypes matched during conversion, and the biotypes of the unmatched transcripts (**Figure 1B**). The system generates a summary table displaying both successfully matched genes and those without positive matches. This table is available to download as a .CSV file from within the application.

Importantly, not all mouse transcripts have human orthologs, owing to genomic and species-level differences in post-translational modifications, alternative RNA splicing, etc. During the process of conversion of mouse transcripts (or any other species) to human transcripts, there will inevitably be some measurable loss of information – genes or transcripts that are unique to the mouse transcriptome, with no annotated human orthologs, will not be available for downstream analysis after conversion. The Ensembl BioMart DB places a natural limit on OrthologAL’s capacity, thus the conversion efficiency is dependent on the input dataset itself, and the transcripts detected within it.

To make comparisons across datasets and disease types as consistent as possible, we measure “conversion efficiency” as the proportion of HGNC gene symbol counts after conversion to the total number of species-level gene symbols profiled in the dataset. We further quantify data “loss” by also calculating the “protein-coding conversion efficiency” as the ratio of protein-coding HGNC symbols post-conversion to the number of protein-coding gene symbols before conversion. protein-coding genes are also much more deeply annotated in the BioMart DB than other transcript biotypes such as microRNAs (miRNAs) or small nuclear RNAs (snRNAs), which also allows for better comparisons between species and data types.^26,27^

To assess the data efficiency of OrthologAL in the context of shared-nearest-neighbor (SNN) clustering^28^, we applied parallel clustering to both the original dataset expression assay and the OrthologAL converted expression assay. Raw counts or UMIs were log-normalized and scaled using Seurat. SNN clustering was performed in Seurat utilizing matching resolution parameters. Using the two sets of annotated data (raw and converted), the RAND index^29^ was calculated to determine the efficiency of SNN clustering to assign cells the same cluster identities after OrthologAL-mediated conversion. A calculated RAND index varies between 0 and 1, where 1 signifies a complete agreement between clustering for all cells, and A RAND value of 0 indicates the clustering does not agree for any cells (**Figure 1B**).^30^

To assess the data efficiency of OrthologAL in the context of discrete cell types within scRNA-seq and snRNA-seq datasets, we quantified the data “lost” within each cell due to conversion as a measure of the percentage of counts (or UMIs) retained within a cell in the converted assay relative to the counts per cell present within the original assay. This is then visualized on a per-cluster or per-cell-type annotation basis using violin plots (**Figure 1B**).

OrthologAL also takes spatialRNA-seq datasets as input. Given inherent differences in capture approaches between these and sc- or snRNA-seq data, we quantified data “lost” for spatialRNA-seq data on a “per-pixel” basis (**Figure 1B**). Specifically in the use-case presented, we quantify per-pixel retention in 10X Visium^31,32^ data of an MB PDOX mouse model across 4 images as a proportion of counts or UMIs retained within the humanized expression matrix from the original mm10-aligned (mus musculus) expression data.

## 3 RESULTS

**OrthologAL facilitates the humanization of non-human gene expression data for single-cell, single-nuclei, and spatial RNA sequencing formats.**

OrthologAL is available as an R package at www.github.com/AyadLab/OrthologAL. Once installed and loaded, the OrthologAL Shiny app can be launched with the single command: *OrthologAL::RunOrthologAL()*. To benchmark the efficacy and measure QC of the humanization and harmonization of pre-clinical high-resolution transcriptomics data, we applied OrthologAL to single-cell, spatial, and single-nuclei transcriptomic datasets from mouse models of medulloblastoma (MB) and both mouse and rat models of spinal cord injury (SCI). These use cases also serve to test whether OrthologAL can handle orthologous gene conversion from different species, medical conditions, and animal models across distinct technologies.

### 3.1 USE CASE 1: Medulloblastoma

MB is the most common pediatric CNS malignancy. To demonstrate the utility of OrthologAL on preclinical models of MB, we applied it to publicly available scRNA-seq data from a GEMM of SHH MB (C57BL/6 SmoM2-eYFP^loxP/loxP^)^24^. To examine the application of OrthologAL to pixel-based spatialRNA-seq data for harmonization of mixed mouse and human transcripts, we applied it to 10X Visium data of a PDOX model of SHH MB.

The BioMart DB contains information on 55,414 mouse Ensembl IDs, of which 53,041 are unique MGI-annotated transcripts. Of 55,414 Ensembl IDs, 22,563 are protein-coding. From 53,041 MGI-annotated transcripts, 21,884 are unique protein-coding transcripts for the mouse (**Extended Figure 1A**). In total, 18,776 HGNC-annotated genes (of which 17,620 are protein-coding) (**Extended Figure 1B**) are matched orthologs between humans and mice in BioMart. Therefore, 80.6% (17,620/21,884) of mouse protein-coding MGI genes have a matched human ortholog. This represents the efficiency limit for converting a mouse gene expression dataset to that of protein-coding human orthologs.

We successfully used OrthologAL to convert the SmoM2 model (GSE129730)^24^ scRNA-seq dataset from mouse to human orthologs. The SmoM2 dataset contains expression data for 19,925 unique murine Ensembl IDs representing 19,905 unique MGI-annotated genes, of which 14,698 are protein-coding transcripts (**Figure 2A**). After conversion with OrthologAL, we identified 14,298 unique HGNC-annotated human orthologs, of which 14,058 are protein-coding (**Figure 2A**). Importantly, these results highlight that the majority of documented human orthologs in the BioMart DB are protein-coding genes. Therefore, out of the total number of MGI protein-coding genes present in the SmoM2 scRNA-seq data, 95.6% of them had human orthologs. In total, the dataset contained 79.78% (14,058 / 17,620) of all documented human orthologs (**Figure 2B**).

**Figure 2.**
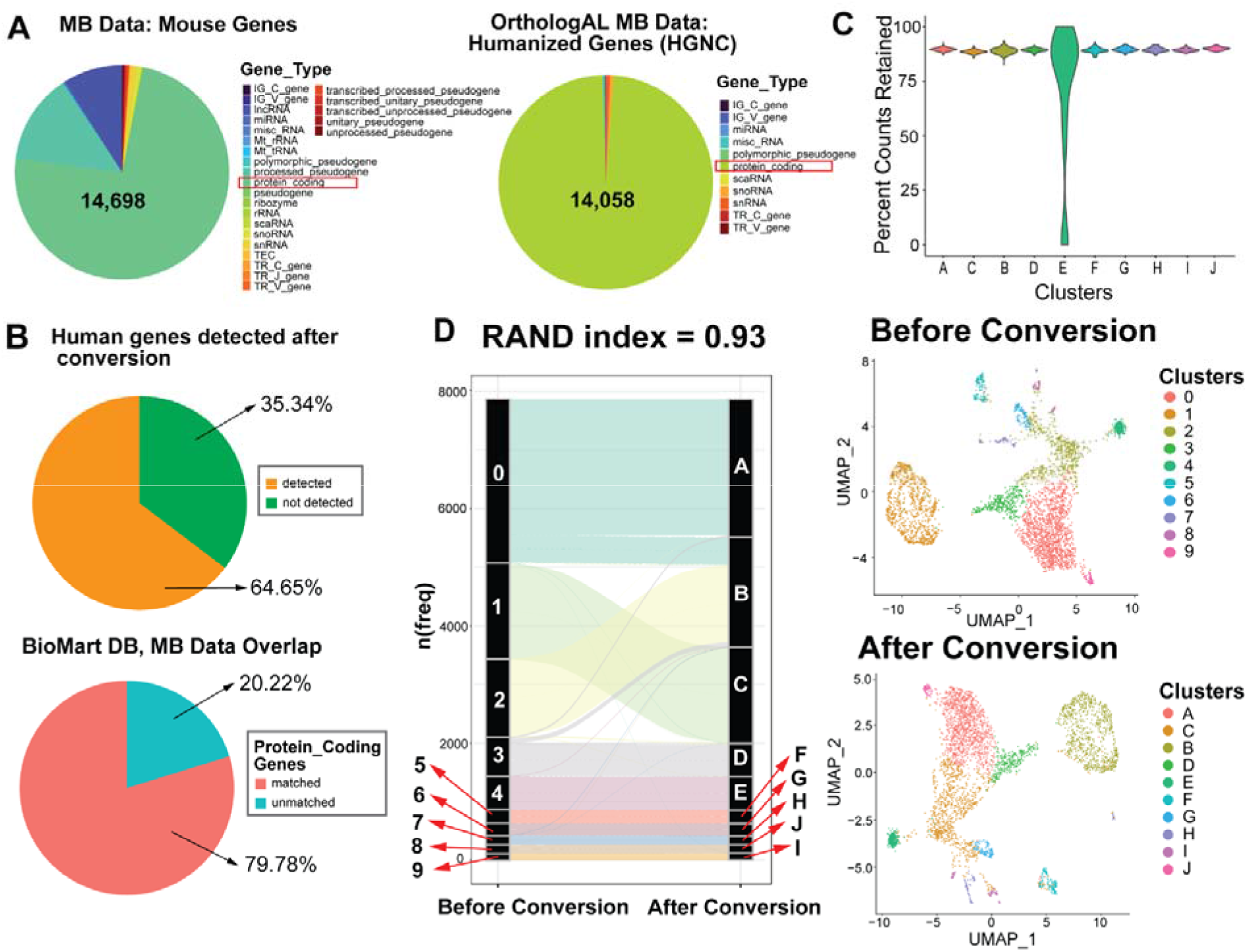
OrthologAL is a data-efficient method for the humanization of mouse scRNA-seq data. **A)** Distribution of transcripts present in the mouse MB GEMM (C57BL/6 SmoM2-eYFP^loxP/loxP^, Ocasio et al., 2019) scRNA-seq dataset (left) and distribution of genes present after orthologous conversion using OrthologAL (right). protein-coding subset is highlighted by a red box and quantified in the respective pie charts. **B)** Pie chart showing the proportion of genes with counts greater than zero that were successfully converted to human orthologs (top); Pie chart showing the proportion of human orthologous genes profiled by the mouse MB scRNA-seq dataset (bottom).**C)** Violin plot showing the proportion of gene counts per cell retained after orthologous conversion, stratified by SNN cluster identity. **D)** Alluvial plot showing cell assignments to SNN clusters before and after conversion. SNN clusters are labeled from 0-9 on the left (Before Conversion) and A-J on the right (After Conversion). UMAP plots before (top) and after (bottom) conversion. Cluster labels are named to match the alluvial plot.

Importantly, we also quantified conversion efficiency specifically for detected genes, regardless of their protein-coding status. The total number of MGI genes detected before conversion, with UMI counts greater than zero, was 21,948. After conversion, the number of HGNC-annotated human genes with counts greater than zero in the data table was 14,191. Therefore, 64.65% of genes detected within this dataset were retained after conversion (**Figure 2B**). In contrast to comparing only protein-coding genes, this lower percentage emphasizes that the majority of those genes lost to orthologous conversion are pseudogenes, alternative splicing variants, and other non-coding RNAs.

After conversion, we wanted to validate the preservation of important transcriptomic information that could potentially be lost by analyzing only orthologous genes. Using both the original SmoM2 mouse-aligned expression data and the data retained following humanization, we performed parallel SNN clustering to confirm that similar clusters are identified after conversion. For quantitative validation, we calculated the RAND index utilizing these cluster identities. The RAND index measures the similarity of clustering results before and after conversion with a perfect alignment, giving a RAND index of 1. The resulting uniform manifold approximation projection (UMAP) from SNN and subsequent dimensional reduction from both before and after conversion are shown in Figure 2D and visually demonstrate a high degree of cluster retention after ortholog conversion. The flow of single-cell assignments to clusters before and after conversion is shown in the alluvial plot^33^ (**Figure 2D**), which confirms this high degree of retention of cluster identity. Importantly, the cluster similarity on the SmoM2 dataset after humanization had a high agreement score with a RAND index of 0.93 (93% agreement).

As another measure of data preservation by OrthologAL-mediated conversion, and to determine any cell-type specific patterns of information loss, we quantified the proportion of counts retained within single cells after conversion relative to the counts per cell present within the original assay. The results of this analysis are shown in Figure 2C, with approximately 85% of counts being retained among single-cells on a SNN cluster level. Counts lost represent those associated with genes without annotated human orthologs.

Next, we sought to apply OrthologAL to spatialRNA-seq data. Researchers often use PDOX models to study MB and other tumors. As a result, spatial sections will inevitably contain both mouse (normal brain) and human (PDOX) genetic material. To account for this, we designed OrthologAL with a specific PDOX setting to be able to recognize and convert expression data that contains both mouse and human transcripts. We acquired 10X Visium data from a PDOX model of SHH MB from Vo et al.^23^ and humanized the dataset using OrthologAL, applying the aforementioned PDOX settings.

The original SHH MB PDOX spatialRNA-seq data is comprised of 50.6% mm10-aligned (mouse) and 49.4% hg38-aligned (human) expression (**Figure 3C**). In total, the PDOX spatialRNA-seq data contains transcriptomic expression information on 75,392 unique Ensembl IDs, consisting of both mouse and human transcripts. Of these, 42,410 are unique protein-coding transcripts (21,174 mouse and 21,236 human). Before conversion, there were 39,990 unique protein-coding MGI- or HGNC-annotated transcripts. Of these, 21,153 represent unique protein-coding MGI-annotated genes.

**Figure 3.**
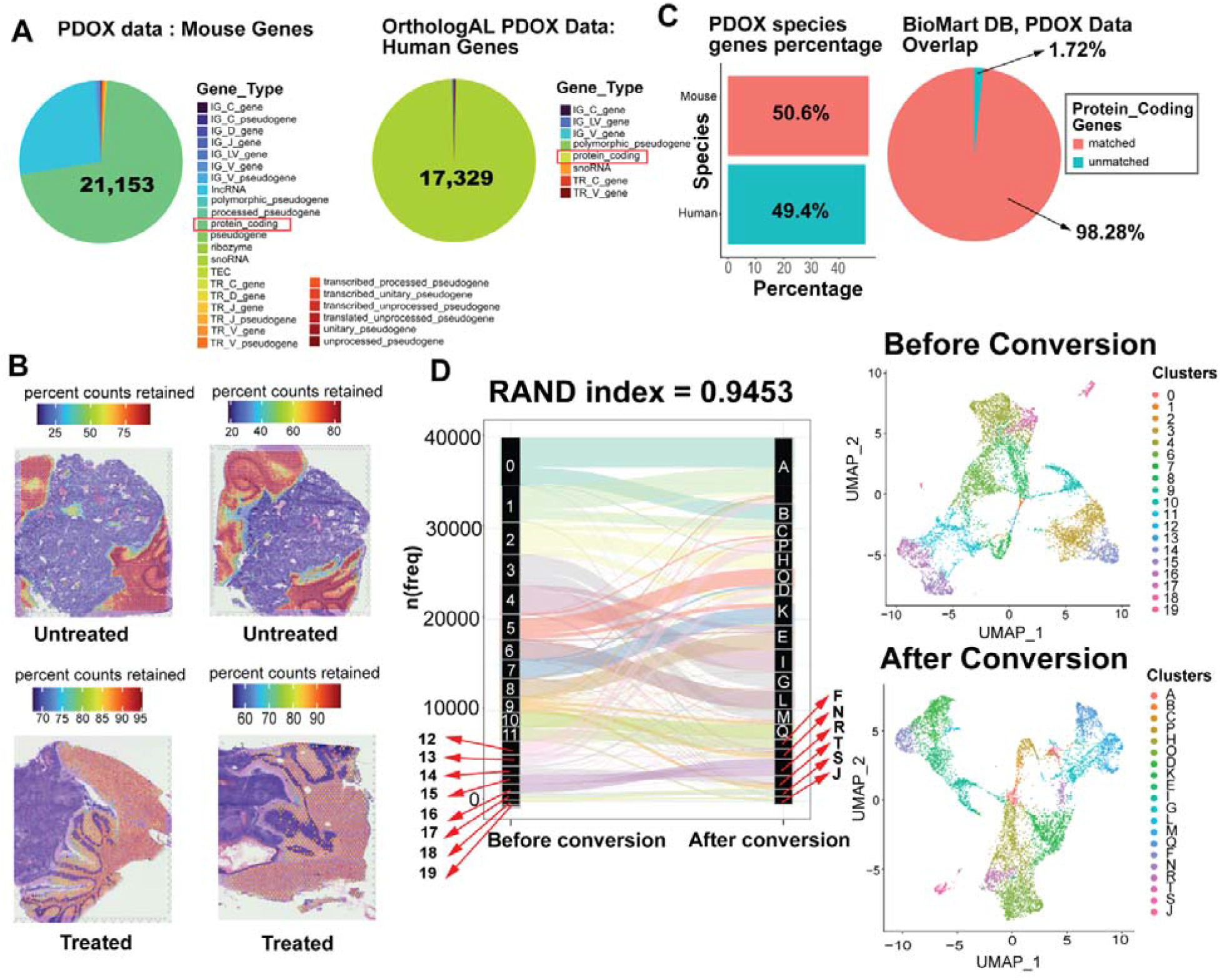
OrthologAL yields a high RAND index after the conversion of non-human genes to human genes in spatial transcriptomic data. **A)** Distribution of genes present in the mouse MB PDOX spatialRNA-seq dataset (left) and distribution of genes present after orthologous conversion using OrthologAL (right). protein-coding subset is highlighted by a red box and quantified in the respective pie charts. **B)** Spatial feature plots showing the percentage of data retained per spot/pixel after orthologous conversion. Percent retained is calculated for each pixel containing mouse cells as the ratio of converted counts retained in the ortholog-converted expression data relative to the total counts present for the same pixel in the original mm10 expression data. **C)** Bar plot demonstrating the proportion of mouse and human genes present in the MB PDOX spatialRNA-seq dataset (left). Pie chart showing the proportion of human orthologous genes profiled by the mouse MB PDOX dataset (right). **D)** Alluvial plot showing cell assignments to SNN clusters before and after conversion. SNN clusters are labeled from 0-19 on the left (Before Conversion) and A-T on the right (After Conversion). UMAP plots before (top) and after (bottom) conversion. Labels correspond to identities annotated in the alluvial plot in D.

Following conversion with OrthologAL, we identified 17,329 unique HGNC-annotated human orthologs matched to the original mm10 expression. (**Figure 3A**). As with the MB scRNA-seq data, we performed dimensional reduction and SNN clustering on the data before and after conversion, showing conservation of cluster identity following ortholog conversion. (**Figure 3D**). Quantitatively, the spatialRNA-seq data before and after conversion have a high agreement score of 94.53% (0.9453 RAND index). We also evaluated the percentage of gene counts retained after conversion on a spot-by-spot (pixel-by-pixel) basis. To do this, we calculate the percent counts retained at each spot with mm10 detection as the total protein-coding HGNC-converted gene counts at the spot after conversion divided by the total number of mm10 gene counts for the spot before conversion. Though we show high percent counts retained across most spots, we observe the loss of counts near the tumor/brain border, which we attribute to the detection of both human and mouse cells and therefore genes at these spots (**Figure 3B**).

### 3.2 USE CASE 2 : Spinal Cord Injury (SCI)

Spinal cord injury (SCI) is a neurological condition caused by damage to the spinal cord, typically the result of a sudden traumatic injury from a motor vehicle accident or a fall.^20,34^ We acquired a scRNA-seq dataset from a rat model of SCI from Li et al. 2022 (GSE213240)^35^ as another species test for OrthologAL implementation. Tabulae Paralytica from Skinnider et al. 2023^36^ is a database that compiles multiple SCI atlases and contains snRNA-seq data of half a million spinal cord cells from mouse models of SCI. We applied OrthologAL to both of these datasets. Importantly, these represent both different animal models (rat and mouse) and different data types (scRNA-seq and snRNA-seq) in the same setting.

The rat species (*rattus norvegicus*) has 30,560 total Ensembl-annotated transcripts in BioMart. Of these, 23,096 are unique protein-coding transcripts and 17,766 are unique protein-coding RGD-annotated (Rat Genome Database) transcripts (**Extended Figure 1C**). Importantly, 16,684 of the protein-coding RGD-annotated genes have HGNC-annotated human orthologs in BioMart DB.

The SCI rat scRNA-seq dataset contains 8 rat samples with SCI classified from mild to severe. It contains 20,236 total unique Ensembl-annotated transcripts, of which 19,305 are protein-coding that map to 19,273 unique protein-coding RGD-annotated gene symbols (**Figure 4A**). OrthologAL-mediated conversion from rat to human resulted in 16,328 unique HGNC-annotated genes, of which 16,082 were protein-coding (**Figure 4A**). Importantly, of the 16,684 total possible HGNC human-rat orthologs, 96.39% were profiled in this dataset. Additionally, 90.52% of protein-coding RGD-annotated genes were successfully converted to orthologs in this dataset (**Figure 4B**). Prior to conversion, there were 20,236 RGD-annotated genes with counts greater than zero, and after there were 15,522 HGNC-annotated genes that met these criteria, meaning 76.71% of gene counts were retained by conversion (**Figure 4B**).

**Figure 4.**
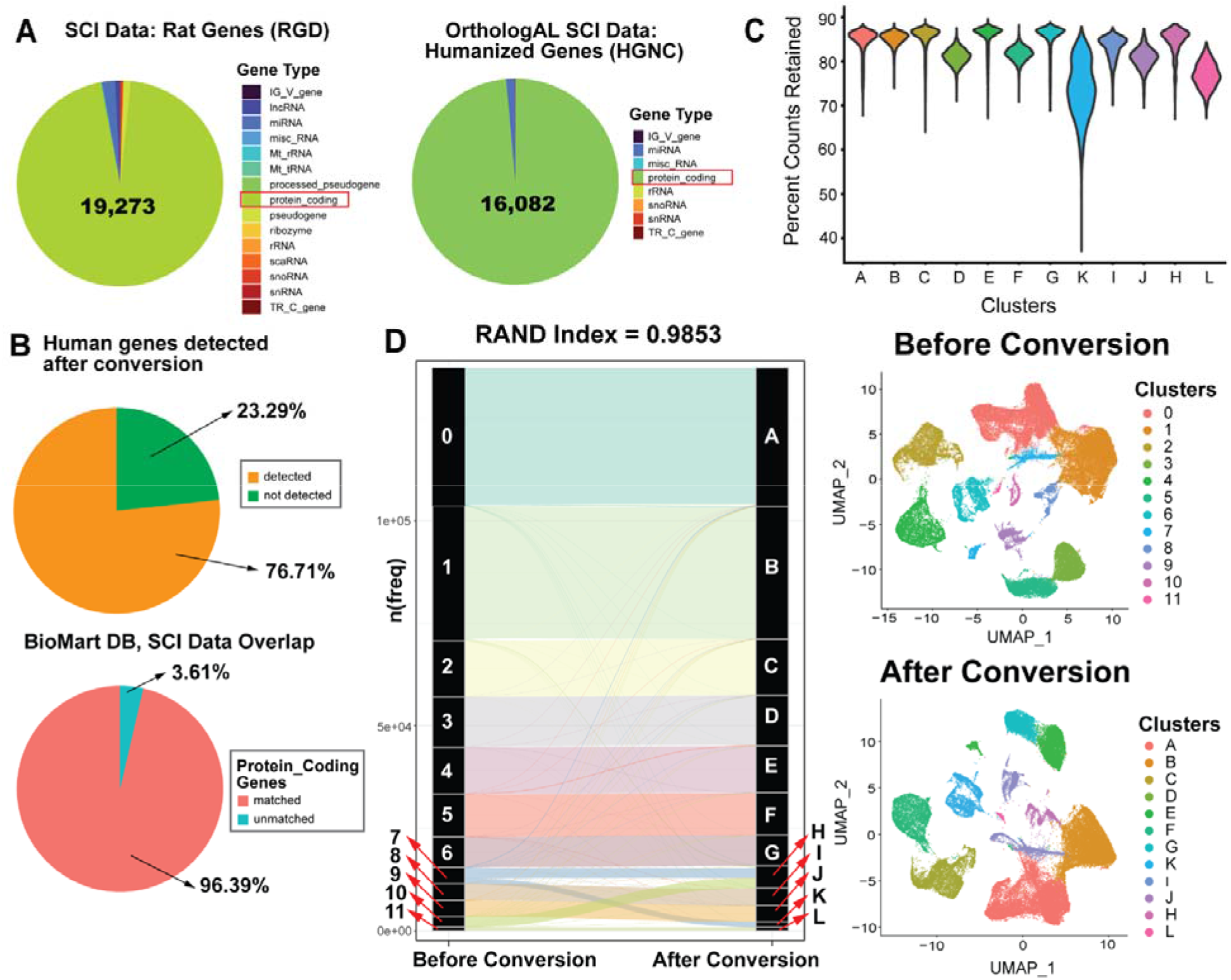
OrthologAL yields a high RAND index for conversion of rat genes to human orthologs in single-cell RNA-seq data. **A)** Distribution of transcripts present in the SCI rat scRNA-seq dataset (left) and distribution of genes present after orthologous conversion using OrthologAL (right). protein-coding subset is highlighted by a red box and quantified in the respective pie charts. **B)** Pie chart showing the proportion of genes with counts greater than zero that were successfully converted to human orthologs (top); Pie chart showing the proportion of human orthologous genes profiled by the rat SCI dataset (bottom). **C)** Violin plot showing the proportion of gene counts per cell retained after orthologous conversion, stratified by SNN cluster identity. **D)** Alluvial plot showing cell assignments to SNN clusters before and after conversion. SNN clusters are labeled from 0-11 on the left (Before Conversion) and A-L on the right (After Conversion). UMAP plots before (top) and after (bottom) conversion. Cluster labels are named to match the alluvial plot.

As another measure of data preservation by OrthologAL-mediated conversion, we quantified the proportion of counts retained within single cells after conversion relative to the counts per cell present within the original assay. The results of this analysis are shown in Figure 2C, with approximately 85% of counts being retained among single cells across all SNN clusters.

Through analysis of SNN clusters before and after conversion, we found that we achieved 98.53% cluster similarity (RAND score of 0.9853) with this dataset by using OrthologAL (**Figure 4D**). Moreover, this high concordance rate is exemplified by the UMAP plots in (**Figure 4D**) that visually show high conservation of clusters and dimensional architecture.

The Tabulae Paralytica snRNA-seq dataset contains expression data on 30,684 unique Ensembl IDs and 30,643 unique MGI-annotated genes of which 21,424 are mouse protein-coding genes. OrthologAL was able to convert 17,609 total MGI-annotated genes to HGNC symbols, of which 17,485 of these are protein-coding orthologs. Out of 17,620 total protein-coding mouse-human orthologs in the BioMart database, the dataset contained 99.23% of orthologous genes (**Figure 5B**). Of 26,918 total MGI-annotated genes with counts greater than zero, 17,422 were successfully converted to human orthologs with non-zero counts (64.72%, **Figure 5B**).

**Figure 5.**
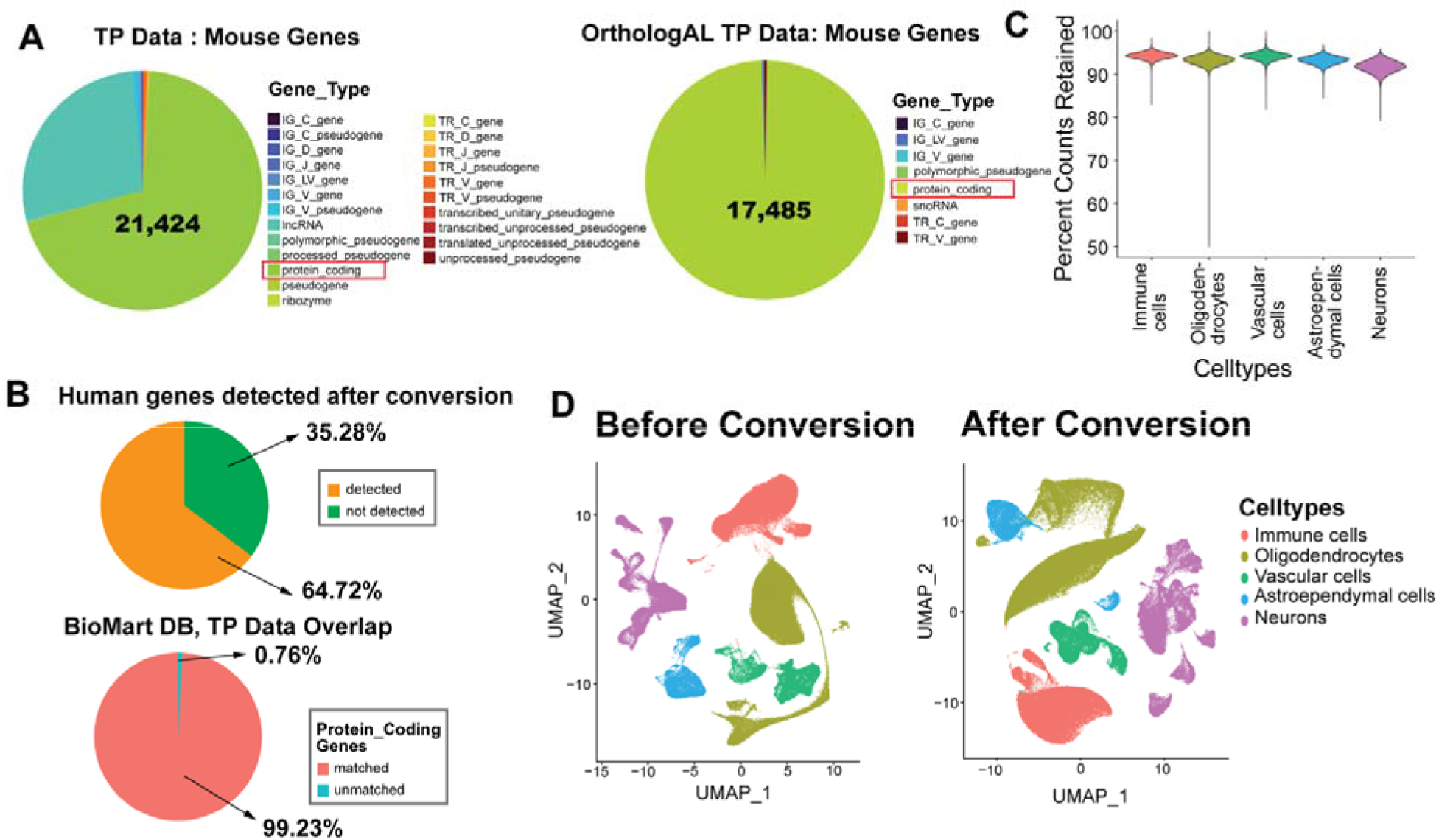
OrthologAL app applied to Tabula Paralytica SCI single-nuclei data shows high concordance after conversion. **A)** Distribution of transcripts present in the Tabulae Paralytica dataset (left) and distribution of genes present after orthologous conversion using OrthologAL (right). protein-coding subset is highlighted by a red box and quantified in the respective pie charts. **B)** Violin plot showing the proportion of gene counts per cell retained after orthologous conversion, stratified by cell type. **C)** Pie chart showing the proportion of genes with counts greater than zero that were successfully converted to human orthologs (top); Pie chart showing the proportion of human orthologous genes profiled by the Tabulae Paralytica dataset (bottom). **D)** UMAP plots before (left) and after (right) conversion, colored by cell type annotation.

The Tabulae Paralytica dataset consists of multiple layers of cell type annotations ranging from broad cell classifications to specific cell lineages, which were annotated using an automated approach^37^. We observed that cell types such as oligodendrocytes, neurons, and immune cells have transcript counts retained at a high rate, approximately 90-95% of transcripts retained after human conversion (**Figure 5C**). Furthermore, through analysis of SNN clusters before and after conversion, we found that we achieved 93.55% cluster similarity (RAND score of 0.9355) with this dataset after using OrthologAL (**Figure 5D**).

## 4 DISCUSSION

An emerging method in drug discovery is disease signature reversal, whereby the transcriptional profile of one sample from a “disease state” is compared to that of the corresponding normal tissue ^16,38,39^. Large pharmacotranscriptomic datasets used in such methods are generated from human cell lines, but GEMMs and other animal models are often necessary to study complex conditions such as MB and SCI. This makes it challenging to apply disease signature reversal and other drug discovery methods to animal model datasets. Therefore, a simple-to-use method is needed to convert non-human transcriptomic data to that of the corresponding human orthologs and facilitate direct comparison using pharmacogenomic or pharmacotranscriptomic databases.

Collectively, our findings demonstrate that OrthologAL performs exceptionally well in converting non-human species genes to human orthologs in two use cases across three different data types. In each species, condition, and data type, the RAND index was always greater than 0.9, which suggests that our conversion method is robust. Additionally, the OrthologAL app makes Ensembl data easily accessible to researchers eliminating the data wrangling typically required to reformat a gene expression dataset for use with BioMart. Importantly, OrthologAL primarily takes Seurat objects as input, a novel feature that makes orthologous gene conversion for high-dimensional single-cell and spatial transcriptomic data a simple matter of file upload. Compatibility with the most popular single-cell data analysis R tool also allows OrthologAL to calculate and present the user with single-cell QC metrics and preserve the original data in a separate assay in the same Seurat object. Researchers can easily install the OrthologAL package in R with minimal coding experience^40^.

OrthologAL has multiple applications and advantages over existing options, especially the ease of use and access. While existing tools such as syngo portal, and the R packages orthogene and iGEAK do facilitate the conversion of non-human genes to their human orthologs using HUGO and HGNC data, they take expression matrices or gene lists as input and do not generate quality control metrics.^41–43^ That OrthologAL takes a Seurat object as input, means that it can handle diverse data types including spatial transcriptomics datasets. Further, OrthologAL coerces this output into a downloadable Seurat object, and is capable of handling dual-species input such as that found within PDOX datasets..

With the emergence of high-dimensional single-cell technologies, tools that are able to convert single-cell genomic and transcriptomic data while preserving them in their main analytic environments (Seurat objects) are critical. Users are able to run OrthologAL using their preference of scRNA-seq, snRNA-seq, or spatialRNA-seq data from non-human species, or from datasets that contain mixed-species gene expression. OrthologAL produces simple-to-understand quality control metrics, and allows for the easy identification of genes which were successfully mapped to the BioMart DB. These quality control metrics are adapted to both single-cell resolution and spatial data. Importantly, we have shown that OrthologAL performs well on modern transcriptomic technologies including scRNAseq, snRNA-seq and spatialRNA-seq. That this method works well in mixed spatial PDOX datasets emphasizes its functionality. Further modifications and improvements of OrthologAL will enhance its utility and ease of use in the scientific and medical communities. Ultimately, this tool will be essential in research pipelines working toward the repurposing and discovery of new therapeutics for human disease.

## DATA AVAILABILITY

OrthologAL is available for download as an R package with functions to launch the Shiny GUI at https://github.com/AyadLab/OrthologAL.

The single-cell Medulloblastoma data was downloaded from the NCBI Gene Expression Omnibus with the identifier GSE129730, and is available at https://www.ncbi.nlm.nih.gov/geo/query/acc.cgi?acc=GSE129730.

10X Visium of PDOX models from Vo et al., was acquired through contacting the authors. The spatial transcriptomics raw data has been made available and can be downloaded from ArrayExpress under the link https://www.ebi.ac.uk/biostudies/arrayexpress/studies/E-MTAB-11720.

The single-cell and single-nuclei rat spinal-cord injury raw data are deposited in the Gene expression Omnibus under GSE213240 and GSE234774 and can be found at https://www.ncbi.nlm.nih.gov/geo/query/acc.cgi?acc=GSE213240 and https://www.ncbi.nlm.nih.gov/geo/query/acc.cgi?acc=GSE234774 respectively.

## ACKNOWLEDGEMENTS

We thank all members of the Ayad, Jae Lee, and KiBum Lee laboratories for fruitful discussions. This work was supported by RM1NS133003 and NS118023. We thank the laboratories of Quan Ngyugen and Laura A Genovesi for their support in providing access to their 10X Visium data.

