## Supplement Figure 1 for "OrthologAL: A Shiny application for quality-aware humanization of non-human pre-clinical high-dimensional gene expression data"

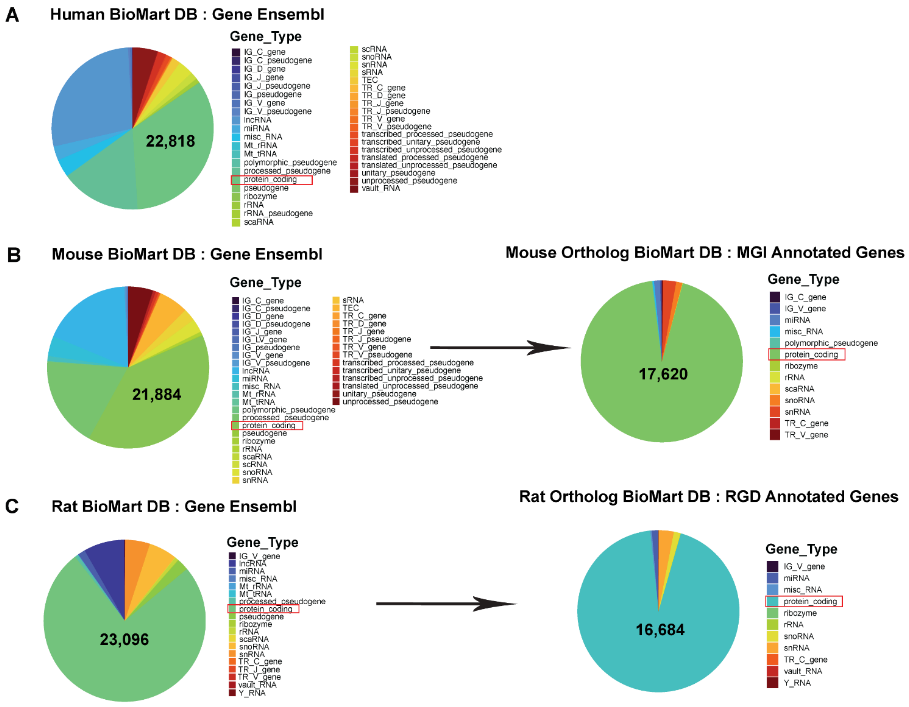


**Extended Figure 1. OrthologAL facilitates the conversion of available species data from BioMArt Database to human genes.**

**A)** Distribution of human transcripts present in the BioMart DB and protein coding genes are highlighted by a red box **B)** Distribution of mouse transcripts present in the BioMart (left) and distribution of orthologous conversion of the whole rat database (right) where protein coding genes are highlighted by a red box. **C)** Distribution of rat transcripts present in the BioMart (left) and distribution of orthologous conversion of the whole rat database (right) where protein coding genes are highlighted by a red box.
